# Dietary potassium and cold acclimation additively increase cold tolerance in *Drosophila melanogaster*

**DOI:** 10.1101/2024.05.24.595710

**Authors:** Bassam Helou, Marshall W. Ritchie, Heath A. MacMillan, Mads Kuhlmann Andersen

**Author notes:** Corresponding author: Mads Kuhlmann Andersen.

## Abstract

In the cold, chill susceptible insects lose the ability to regulate ionic and osmotic gradients. This leads to hemolymph hyperkalemia that drives a debilitating loss of cell membrane polarization, triggering cell death pathways and causing organismal injury. Biotic and abiotic factors can modulate insect cold tolerance by impacting the ability to mitigate or prevent this cascade of events. In the present study, we test the combined and isolated effects of dietary manipulations and thermal acclimation on cold tolerance in fruit flies. Specifically, we acclimated adult *Drosophila melanogaster* to 15 or 25°C and fed them either a K^+^-loaded diet or a control diet. We then tested the ability of these flies to recover from and survive a cold exposure, as well as their capacity to protect transmembrane K^+^ gradients, and intracellular Na^+^ concentration. As predicted, cold-exposed flies experienced hemolymph hyperkalemia and cold-acclimated flies had improved cold tolerance due to an improved maintenance of the hemolymph K^+^ concentration at low temperature. Feeding on a high-K^+^ diet improved cold tolerance additively, but paradoxically reduced the ability to maintain extracellular K^+^ concentrations. Cold-acclimation and K^+^-feeding additively increased the intracellular K^+^ concentration, aiding in maintenance of the transmembrane K^+^ gradient during cold exposure despite cold-induced hemolymph hyperkalemia. There was no effect of acclimation of diet on intracellular Na^+^ concentration. These findings suggest intracellular K^+^ loading and reduced muscle membrane K^+^ sensitivity as mechanisms through which cold-acclimated and K^+^-fed flies are able to tolerate hemolymph hyperkalemia.

**Highlights:**

- Insect cold tolerance varies in relation to ionoregulatory capacity
- Cold acclimation improves cold tolerance and K^+^ handling during cold exposure
- A high K^+^ diet also improves cold tolerance, but reduces the K^+^-handling capacity
- We highlight a novel mechanism for preventing K^+^ gradient disruption

## Introduction

Insect physiology is strongly influenced by the thermal environment, and their ability to tolerate thermal extremes dictates their potential thermal niche. Cold tolerance, in particular, is a strong predictor of insect latitudinal distribution (Addo-Bediako et al., 2000; Denlinger and Lee, 2010; Overgaard et al., 2011; Kellermann et al., 2012; Lee, 2012; Lancaster, 2016), so understanding the physiological mechanisms that underlie cold tolerance limits and variation therein is important if we want to understand or predict potential range shifts in a changing environment.

Some insects can tolerate freezing of their extracellular fluids (‘freeze tolerant’) while others avoid freezing altogether by lowering the melting point of their bodily fluids (‘freeze avoiding’). Most insects, however, are unable to tolerate temperatures above those that cause freezing and succumb to the effects of cooling on their physiology (Salt, 1961; Lee, 1991; Bale, 2002; Sinclair et al., 2003). When exposed to stressful cold, these ‘chill susceptible’ insects enter a state of neuromuscular paralysis called chill coma (Hazell and Bale, 2011; MacMillan and Sinclair, 2011a). Chill coma is initiated by a rapid loss of nervous system function culminating in a complete shutdown of the CNS (Staszak and Mutchmor, 1973; Robertson et al., 2017; Andersen et al., 2018; Robertson et al., 2020; Andersen et al., 2023; Robertson et al., 2023). Interestingly, chill coma is reversible and if an insect is brought back to permissive temperatures shortly after losing function it will recovery all bodily function seemingly without harm (Overgaard and MacMillan, 2017). By contrast, exposure to severe or prolonged stressful cold results in a systemic collapse of ion gradients, where a disruption of Na^+^ and water balance leads to extracellular hyperkalemia (elevated K^+^ concentration), which, combined with the direct effect of low temperature, depolarizes excitable tissues and worsens the coma (MacMillan et al., 2014). Thus, in cases of extensive cold exposure, recovery requires recovery of both nervous function and systemic ion gradients, primarily a return from extracellular hyperkalemia to baseline levels (MacMillan and Sinclair, 2011b; Andersen and Overgaard, 2019). Despite the possibility of recovery, most chill susceptible insects suffer injury during severe cold exposures, which generally manifest as behavioral deficiencies, locomotor defects, and mortality in severe cases (Overgaard and MacMillan, 2017). Specifically, the combined cellular depolarization caused by cold and extracellular hyperkalemia leads to an intracellular Ca^2+^ overload which triggers cell death pathways (MacMillan et al., 2015c; Bayley et al., 2018; Carrington et al., 2020). Chill susceptible insects that are able to prevent the loss of ion balance at low temperatures have improved cold tolerance and are generally referred to as being more chill tolerant (Koštál et al., 2004; MacMillan et al., 2015a; Andersen et al., 2017c; Overgaard and MacMillan, 2017).

Thermal tolerance limits are plastic and most chill susceptible insects are able to alter their cold tolerance through a variety of processes. Most notable is the ability to increase cold tolerance *via* seasonal or developmental acclimation, which can drastically improve tolerance to chilling (Mellanby, 1954; Andersen et al., 2017a; Weaving et al., 2022). This is also true for the fruit fly *Drosophila melanogaster*, where cold acclimation has been shown to promote adaptive changes in commonly-used chill tolerance phenotypes (Sinclair et al., 2015): 1) chill coma onset temperature (MacMillan et al., 2015b; Schou et al., 2017), 2) chill coma recovery times (Ayrinhac et al., 2004; Ransberry et al., 2011; MacMillan et al., 2015b), and survival (Overgaard et al., 2008; MacMillan et al., 2017b). Central to these improvements in chill tolerance is an enhanced ability to maintain ion balance or better tolerate its collapse without suffering substantial tissue damage. In the hemolymph this is achieved by the combined actions of the Malpighian tubules and the hindgut, which, through a cycle of selective secretion and reabsorption, maintain hemolymph volume and ion concentrations (Edney, 1977; Phillips, 1981). Improvements to insect cold tolerance have repeatedly been linked to the capacity for each of these organs to maintain their osmoregulatory function at low temperatures and thereby prevent hyperkalemia. Specifically, more cold tolerant con- or allospecifics have Malpighian tubules that better maintain overall transport rates and selective K^+^ secretion (MacMillan et al., 2015a; Yerushalmi et al., 2018) and hindguts that better reabsorb water and Na^+^ and selectively reabsorb less K^+^ at low temperature (Andersen and Overgaard, 2020). Combined with improved epithelial barrier function and reduced ion leakage (Andersen et al., 2017c; MacMillan et al., 2017b) these chill tolerant species are able to mitigate or prevent hemolymph hyperkalemia during cold stress (see the review by Overgaard et al. (2021)).

The ability of the osmoregulatory organs to handle an increased ionic load has also been tested through dietary manipulations, and recent studies have tested whether increased dietary salt load, particularly K^+^, impacts cold tolerance. Feeding activity itself reduces the cold tolerance of migratory locusts (*Locusta migratoria*) with evidence to suggest that this was caused by the increased K^+^ load elicited by the diet (Hoyle, 1954; Andersen et al., 2013). Interestingly, a high-K^+^ diet differentially impacts chill tolerance phenotypes in *D. melanogaster*; it raises the chill coma onset temperature while reducing chill coma recovery times (Yerushalmi et al., 2016). Such effects of dietary manipulations also extend to other ions: In crickets, a high Na^+^, rather than K^+^ diet, speeds chill coma recovery but worsens the debilitating effects of prolonged cold stress (Lebenzon et al., 2020). While the effects of these ions varies depending on the species and cold tolerance metric used, improved chill coma recovery times on a high K^+^ diet are supported by earlier observations of changes to the Malpighian tubules, and part of the gut, that counteract the increased ion load by keeping hemolymph K^+^ balanced (Naikkhwah and O’Donnell, 2011; Naikkhwah and O’Donnell, 2012). These salt-induced improvements to renal function are similar to those associated with improved chilling tolerance on standard diets (see Overgaard et al. (2021)), which raises the question of whether there is overlap in the physiological mechanism(s) underlying dietary salt and cold tolerance.

In the present study we sought to investigate how the isolated and combined effects of an increased dietary K^+^ load and cold acclimation improved the cold tolerance of *D. melanogaster*. We first repeated a previous experiment on the effect of dietary salt stress on cold tolerance. We then used a fully-crossed experimental design to examine how increased dietary K^+^ load and cold acclimation affected cold tolerance and the ability to maintain K^+^ balance during stressful cold exposure. We hypothesized that the combination of cold acclimation and a high-K^+^ diet would further increase cold tolerance through an improved ability to handle and prevent increased extracellular K^+^. We could not replicate all earlier observations of the effects of dietary salt stress on cold tolerance, but we did find a strong additive effect of cold acclimation and a high-K^+^ diet on cold tolerance. Furthermore, we found that cold acclimation and a high-K^+^ diet improve tolerance by augmenting the ability to maintain a low extracellular K^+^ concentration during cold exposure and increasing the intracellular K^+^ concentration, effectively mitigating cellular depolarization. Lastly, we posit that the increased intracellular K^+^ concentration observed for K^+^ fed and cold-acclimated flies might act as a secondary K^+^ handling mechanism by which flies store, rather than secrete, excess K^+^ and utilize it to maintain cellular K^+^ gradients (and by extension cell membrane polarization) despite extracellular hyperkalemia.

## Materials and Methods

### Fly rearing and preparation

The population of *Drosophila melanogaster* (Meigen 1830) used in this study was derived from flies collected from London and Niagara on the Lake (Ontatio, Canada) in 2007 (Marshall and Sinclair, 2010). The population was maintained in 200 mL plastic bottles containing ∼50 mL of standard Bloomington food (see Table 1) at 25°C with a 12 h:12h light cycle in a temperature-controlled incubator (MIR-154-PA, Panasonic, Mississauga, ON, Canada).

**Table 1.** Ingredients used for the Bloomington diet (Lakovaara, 1969) and the K^+^- and xylitol-enriched diets (Naikkhwah and O’Donnell, 2011; Yerushalmi et al., 2016). The enriched diet was made by mixing the ingredients in groups A and B separately in 1 L and 250 mL glass beakers respectively, bringing them to a boil and mixing them together under constant stirring. Once the mixture cooled to 55°C, the components in C were added. The mixture was then split into two 500 mL fractions before component D (KCl or xylitol) was added.

| Diet | Ingredients |  | For 1 L of food |
| --- | --- | --- | --- |
| Bloomington | Distilled water<br>Inactive yeast<br>Soy flour<br>Yellow cornmeal<br>Light dry malt extract<br>Agar<br>Corn syrup<br>Propionic acid |  | 920 mL<br>15.9 g<br>9.2 g<br>67.1 g<br>42.3 g<br>8 g<br>70 mL<br>4.4 mL |
| Enriched diet base | A | Tap Water<br>Sucrose<br>Agar<br>KNa-tartrate<br>KH<sub>2</sub>PO<sub>4</sub><br>NaCl<br>MgCl<sub>2</sub><br>CaCl<sub>2</sub> | 800 mL<br>100 g<br>18 g<br>8 g<br>1 g<br>0.5 g<br>0.5 g<br>0.5 g |
|  | B | Tap Water<br>Inactive yeast | 200 mL<br>50 g |
|  | C | Tap Water<br>Propionic acid<br>85% o-phosphoric acid | 5 mL<br>4.55 mL<br>0.45 mL |
| Enriched diet additions | D | KCl<br>or<br>Xylitol | 13.92 g<br>60.86 g |

To generate flies for experiments, adults were transferred into bottles with fresh medium to oviposit for 2 h before being returned. The bottles containing newly-laid eggs were placed back into the 25°C incubator until flies emerged (approximately 10 days). Emerging flies were collected daily over three days and transferred into 40 mL vials containing 10 mL standard Bloomington food. At this time, flies were haphazardly split into two groups for acclimation at either 15°C or 25°C in other incubators (same as above) with the same light cycle. After 2-3 days, female flies were isolated under light CO_2_ anaesthesia and placed into vials containing fresh food. CO_2_ anaesthesia lasted ∼10 min and flies were left to recover for two days afterwards to avoid negative effects on thermal tolerance (Nilson et al., 2006; MacMillan et al., 2017a). After two days, the flies were split again and transferred into 40 mL vials containing 10 mL of one of six different diets (Bloomington, control, and enriched with either sucrose, xylitol, sodium, or potassium; note that the comparison between all diets was only performed on 25°C-acclimated flies) (see **Table 1** for recipes). Of the experimental diets, the control diet formed the base diet into which each of the salts (Na^+^ or K^+^) or osmolites (sucrose or xylitol) were added. Flies were fed on each diet for 24 h prior to experiments to ensure ample consumption (Yerushalmi et al., 2016). The effects of thermal acclimation were only examined in flies fed the potassium- or the xylitol-enriched diet. Here, the xylitol-enriched diet served as an osmotic control to the potassium-enriched diet as xylitol is a non-nutritional carbohydrate that has been used as an osmotic control previously (Lebenzon et al., 2020), and has no adverse effects at our concentration (Choi et al., 2017). Female flies used in experiments were assumed to be non-virgin and were all 5-6 days post-final ecdysis on the day of the experiment.

### Chill coma recovery time

After being fed their respective diets for 24 h, flies were transferred into individual 3.7 mL glass vials with a screw top, assigned numbers that were blind to the experimenter, and placed into a sealed plastic bag. Next, the vials were submerged into an ice-water slurry such that flies were exposed to 0°C for 6 h. At the end of the exposure, the vials with flies were removed from the cold and placed on a lab benchtop at room temperature (22°C) and a stopwatch was started. From here, flies were monitored continuously and deemed to have recovered from chill coma when they regained their righting reflex and became able to stand up, at which point their chill coma recovery time (CCRT) was recorded. Chill coma recovery time was measured in a total of 151 flies in the initial diet experiment, and in a total of 173 flies in the experiment on the interactive effects of a K^+^ diet and acclimation (sees **Table 2** for group-specific sample sizes).

**Table 2.** Sample sizes for datasets describing the cold tolerance of *Drosophila melanogaster* fed either different diets or acclimated to different temperatures while feeding on either a control xylitol diet or a high-potassium diet.

| Experiment | Acclimation | Diet | N |
| --- | --- | --- | --- |
| Six diets' effect on<br>CCRT<br>(Fig. 1A) | 25 | Bloomington | 32 |
|  | 25 | Control | 23 |
|  | 25 | Sucrose | 21 |
|  | 25 | Xylitol | 22 |
|  | 25 | Sodium | 19 |
|  | 25 | Potassium | 21 |
| Acclimation and diet<br>effects on CCRT<br>(Fig. 1B) | 15 | Xylitol | 45 |
|  | 15 | Potassium | 45 |
|  | 25 | Xylitol | 43 |
|  | 25 | Potassium | 40 |
| Acclimation and diet<br>effects on chilling injury<br>(Fig. 1C) | 15 | Xylitol | 102 |
|  | 15 | Potassium | 132 |
|  | 25 | Xylitol | 168 |
|  | 25 | Potassium | 177 |

### Chilling injury

Chilling injury was quantified following a 24 h exposure to 0°C. After being fed their respective diets, flies were transferred into empty, foam-stopped 40 mL plastic vials (10 flies per vial) and submerged in an ice-water slurry for 24 h. Following the cold stress, flies were transferred to new 40 mL vials containing their respective diets and placed back in their respective acclimation conditions (25°C and 15°C) on their side to prevent immobile flies from sticking to the food. After 24 h, each fly was rated on a 4-point scale to estimate the degree of injury sustained: 0) No movement or dead, 1) moving but unable to stand, 2) standing and slight walking, and 3) climbing on walls and walking (El-Saadi et al., 2020). Chilling injury was quantified in a total of 579 flies (see **Table 2** for group-specific sample sizes)

### Extracellular K^+^ concentration

The extracellular K^+^ concentration was estimated by measuring the concentration of K^+^ in the hemolymph using ion-selective microelectrodes as described by MacMillan et al. (2015a).

After feeding on their diets for 24 h, flies were either sampled for their hemolymph directly from their acclimation temperatures or after a 6 h exposure to 0°C (same approach as above but with 20 flies per vial). Hemolymph was extracted *via* antennal ablation under positive air pressure using a custom-built hemolymph extraction device described by MacMillan and Hughson (2014).

Immediately following extraction, the hemolymph was transferred to a dish filled with hydrated paraffin oil and with its bottom covered in a silicone elastomer (Sylgard, Dow Corning Corporation, Midland, MI, USA). The ion-selective microelectrodes used to measure hemolymph [K^+^] were fashioned from glass capillaries (TW150-4, World Precision Instruments, Sarasota, FL, USA) and pulled to a ∼ 1-3 mm tip on a Flaming-Brown micropipette puller (P-1000, Sutter Instruments, Novato, CA, USA). After pulling, the glass microelectrodes were heated to 300°C on a hot plate and silanized in an atmosphere of *N*,*N*-dimethyltrimethylsilylamine (Sigma Aldrich, Saint Louis, MO, USA) under an inverted, heat-resistant glass petri dish. From here the silanized glass microelectrodes were back-filled with a 100 mM KCl solution and front-filled with a K^+^-selective ionophore (K^+^ ionophore I cocktail B, Sigma). Filled electrodes then had their tips rapidly dipped in a solution of poly-vinyl chloride in tetrahydrofuran (10 mg in 3 mL; Sigma) to prevent displacement of the ionophore. Raw voltages from the ion-selective electrode were read by a pH Amp (ADInstruments, Colorado Springs, CO, USA), digitized using a PowerLab A/D converter (3/40, ADInstruments), and fed to a computer running Lab Chart software (version 4, ADInstruments). The circuit was completed, and the samples grounded with a thinly pulled glass electrode (1B200F-4, WPI) filled with a 500 mM KCl solution and connected to ground. Recorded voltages were converted to concentrations by reference to two standards with a ten-fold difference in [K^+^] (10 and 100 mM KCl [LiCl was used to balance total ion activity]) and the following formula:

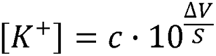

where [*K^+^*] is the concentrations of K^+^ in the hemolymph sample, *c* is the concentrations of K^+^ in the lowest standard solution (here 10 mM KCl), *ΔV* is the voltage difference between the sample and the lowest standard, and *S* is the voltage difference between the lowest and highest concentrations standards (i.e. 10 and 100 mM KCl) referred to as the slope. The slope also served as a quality control step as only electrodes responding to the 10-fold concentration change with a voltage difference close to expected Nernstian potential of 58.2 mV were used (52-60 mV). The slopes of the electrodes used here were 54.0 ± 1.9 mV (mean ± standard deviation). A total of 228 hemolymph samples had their K^+^ concentration measured (see **Table 3** for group-specific samples sizes).

**Table 3.** Sample sizes for experiments measuring extracellular- and intracellular concentrations of K^+^ and Na^+^ of flies acclimated to either 15°C or 25°C and fed either a xylitol control diet or a high-potassium diet, before and after a 6 h exposure to 0°C.

| Ion | Acclimation (°C) | Diet | Exposure time (h) | N<sub>Extracellular</sub> | N<sub>Intracellular</sub> |
| --- | --- | --- | --- | --- | --- |
| $K^+$<br>(Fig. 2) | 15 | Xylitol | 0 | 30 | 9 |
|  | 15 | Xylitol | 6 | 30 | 10 |
|  | 15 | Potassium | 0 | 30 | 10 |
|  | 15 | Potassium | 6 | 33 | 10 |
|  | 25 | Xylitol | 0 | 28 | 8 |
|  | 25 | Xylitol | 6 | 22 | 6 |
|  | 25 | Potassium | 0 | 28 | 8 |
|  | 25 | Potassium | 6 | 27 | 5 |
| $Na^+$<br>(Fig. 3) | 15 | Xylitol | 0 | - | 9 |
|  | 15 | Xylitol | 6 | - | 10 |
|  | 15 | Potassium | 0 | - | 10 |
|  | 15 | Potassium | 6 | - | 10 |
|  | 25 | Xylitol | 0 | - | 8 |
|  | 25 | Xylitol | 6 | - | 6 |
|  | 25 | Potassium | 0 | - | 8 |
|  | 25 | Potassium | 6 | - | 5 |

### Intracellular concentrations of K^+^ and Na^+^

Intracellular concentrations of K^+^ and Na^+^ were estimated by measuring the concentration of both ions in thoracic muscle samples using microwave plasma atomic emission spectrometry (MP-AES).

Flies were fed their diets and sampled at either their acclimation temperature or immediately following a 6 h exposure to 0°C as described above. Thoracic muscle samples were obtained by dry-dissection which involved removing their head, abdomen, legs, wings, and gut after which excess hemolymph was blotted away with a Kim wipe, leaving just the thorax, containing primarily flight muscle. Each sample consisted of four or five thoraces which were placed into pre-weighed, heat-resistant 1.5 mL centrifuge tubes and re-weighed to obtain the wet weight. From here, samples were dried in an oven at 60°C for 24 h after which they were weighed once more on a Sartorius micro balance (ME5, Sartorius Lab Instruments, Goettingen, Germany) to obtain the dry weight which allowed us to calculate the water content for each sample (i.e. wet weight – dry weight). After drying, each sample was fully digested in 220 μL of 70% nitric acid for 2 h at room temperature and 24 h at 60°C, gently mixed on a vortex mixer and spun down in a table-top centrifuge. From each sample, 200 μL was pipetted into a polypropylene test tube (17 mm diameter and 130 mm height, Agilent Technologies, Santa Clara, CA, USA) and diluted to 3.5% nitric acid by adding 3800 μL milliQ water after which all test tubes containing samples were placed into an automatic sampler (SPS 4 Autosampler, Agilent) integrated with the MP-AES (model 4210, Agilent). The MP-AES was allowed to warm up and stabilize for 30 min while continuously aspirating a 3.5% nitric acid solution. Before measuring samples, a standard curve was run with solutions of known K^+^ and Na^+^ concentrations (0.0, 0.1, 0.5, 1.0, and 2.5 mg/L; made by dilution of stock solutions [1000 μg/mL, #5190-8525 and #5190-8503, Agilent]) which allowed us to calculate sample concentrations. A new standard curve was run every 10 samples to account for drift. The MP-AES was programmed to measure both K^+^ and Na^+^ at two wavelengths (K^+^: 766. 491 and 769.897 nm, Na^+^: 588.592 and 588.995 nm) and in triplicates. Negligible differences were found between the concentrations calculated from either wavelength, so their values were averaged and used to calculate concentrations in diluted samples. From this, intracellular concentrations were back-calculated based on the total dilution of the sample water content and converted from mg/L to mmol/L using each element’s molar mass. 66 samples were measured in total on the MP-AES (see **Table 3** for group-specific sample sizes).

### Data analysis

All data and statistical analyses were performed using R software (version 4.2.3; R Core Team (2023)). Before testing for any main effects, all datasets were tested for normality using Shapiro Wilk’s tests (using the shapiro.test() function) and by inspecting boxplots. Many datasets had non-normal distributions with values above the mean being more extreme. In these cases, the datasets first tested for outliers using Rosner’s test using the rosnerTest() function from the *EnvStats* package (Millard, 2013). In some cases this resolved issues with non-normality, however, in cases where it didn’t, a log-transformation resulted in either fully normal-distributed data or a substantial improvement in the normality of the residuals, and we therefore chose to use this transformation and proceed with parametric analyses.

The effect of diet on CCRT was analyzed with a one-way ANOVA, which was followed by a pairwise t tests to identify statistically different diet groups. In this analysis, P values were adjusted using the False Discovery Rate. Furthermore, the Bloomington was excluded in this *post hoc* analysis, as this primarily served to test if there were any effects of switching flies to the control diet (which there wasn’t). The experiment on interactive effects of diet (high K^+^ vs. xylitol-control) and thermal acclimation (15 vs. 25°C) on CCRT were run twice on separate days, and to account for this we analyzed this data with a linear mixed-effect model on log-transformed data using the lme() function from the *nlme* package (Pinheiro, 2009) which included diet and acclimation as fixed factors, and trial as a random factor. The interacting effects of diet and acclimation on chilling injury, as estimated using the scoring system, were also run in two trials and was therefore also analyzed with a linear mixed-effect model, but on non-transformed data.

The effects of acclimation temperature, diet, and cold exposure on the extracellular concentrations of K^+^ were analyzed using a three-way ANOVA on log-transformed data (no three-way interaction was found). Effects of the same experimental design and treatments on the intracellular concentration of K^+^ and Na^+^ were analyzed using the same approach (here too no three-way interaction was found).

Nernst potentials for K^+^ were calculated based on mean values for intra- and extracellular concentrations (measured from different individuals) and therefore only one data point exists for each combination of acclimation, diet, and treatment group. For this calculation, their respective acclimation temperatures or 0°C were used in the equation where appropriate.

The level required for statistical significance was 0.05 in all analyses.

## Results

### Effects of diet and thermal acclimation on fruit fly cold tolerance

In our pilot experiment we fed adult female *D. melanogaster* six different diets to test the impacts of dietary osmolites and electrolytes on their cold tolerance, after which we switched our focus to the interactive effects of a high-K^+^ diet and cold acclimation (**Fig. 1**). In support of earlier reports, we found clear effects of diet on CCRT after a 6 h exposure to 0°C (F_5,132_ = 4.3, P = 0.001) (**Fig. 1A**). Specifically, flies fed the potassium diet recovered fastest at 28.4 ± 1.6 min followed by flies fed the sucrose- (32.5 ± 1.8 min) and xylitol-diets (34.4 ± 1.6 min), with the sodium-, control-, and Bloomington diets leading to the slowest recovery times (38.4 ± 2.4 min, 37.4 ± 2.3 min, and 38.6 ± 1.7 min, respectively). The difference between the potassium- and xylitol diets was only trending towards statistical significance in *post hoc* test (P = 0.109), however, when compared directly (using Student’s t test) the potassium-fed flies recovered faster (t_41_ = −2.5, P = 0.013). Thus, we decided to continue by investigating if the consumption of a diet high in K^+^ affected the way cold acclimation improved cold tolerance, using the xylitol diet as an osmotic control. As expected, we found that cold acclimation improved the CCRT of *D. melanogaster* (F_1,168_ = 118.0, P < 0.001) and cold-acclimated flies recovered ∼ 14 min faster than their warm-acclimated conspecifics on average (**Fig. 1B**). Furthermore, we found here that flies fed a high-K^+^ diet recovered ∼ 7 min faster that those fed the xylitol-enriched control diet (F_1,168_ = 34.0, P < 0.001), while there was no interaction between the high-K^+^ diet and cold-acclimation (F_1,168_ = 0.2, P = 0.628). As expected, these effects of thermal acclimation and diet on CCRT were reflected in the degree of chilling injury sustained after a 24 h exposure to 0°C (**Fig. 1C**): Cold acclimation and diet had additive effects on chilling injury such that cold-acclimated flies sustained less injury and had an average score that was ∼ 1.4 higher than their warm-acclimated conspecifics (F_1,574_ = 183.9, P < 0.001) and flies that were fed the high-K^+^ diet had a score that was ∼ 0.3 higher than those fed the xylitol control diet on average (F_1,574_ = 11.0, P < 0.001). No significant interaction was found between acclimation and diet (F_1,574_ = 1.9, P = 0.166). Specifically, cold-acclimated flies had survival scores of 2.7 ± 0.1 and 2.5 ± 0.1 for high-K^+^ and xylitol diets, respectively, while warm-acclimated flies had scores of 1.4 ± 0.1 and 1.0 ± 0.1.

**Figure 1.**
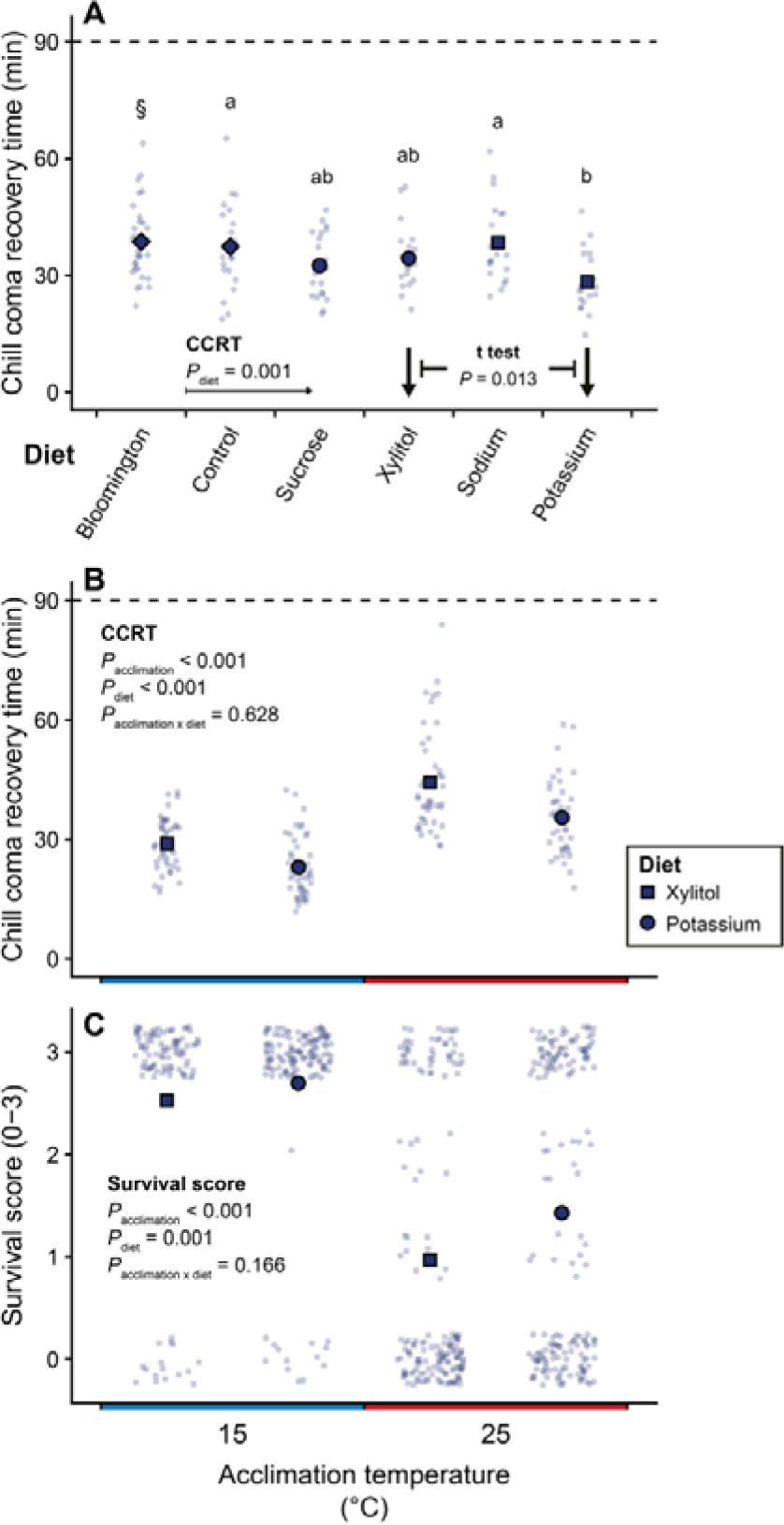
Combined effects of cold-acclimation and a high-K^+^ diet improved the cold tolerance of adult *D. melanogaster*. (A) Chill coma recovery times of *D. melanogaster* following a 6 h exposure to 0°C after having fed on different diets for 24 h (N = 20-32). (B) Chill coma recovery times of cold-exposed *D. melanogaster* acclimated to either 15 or 25°C and fed either a control xylitol diet or a high-K^+^ diet (N = 40-45). (C) Survival scores of *D. melanogaster* exposed to 0°C for 24 h (0-3, higher scores indicate less injury; N = 102-177). Large, opaque points represent mean values and smaller translucent points represent individual data points. In panel A, letters denote statistically different diet groups based on paired t tests (adjusted for false discovery), while the section mark (§) indicates that this group was not included in said test. See **Table 2** for exact, group-specific sample sizes.

### Extra- and intracellular concentrations of K^+^

To investigate whether the observed effects of acclimation and diet related to the ability to maintain transmembrane ion gradients, we measured the extracellular and intracellular concentrations of K^+^, which allowed us to approximate the electrochemical gradient (Nernst potential) (**Fig. 2**).

**Figure 2.**
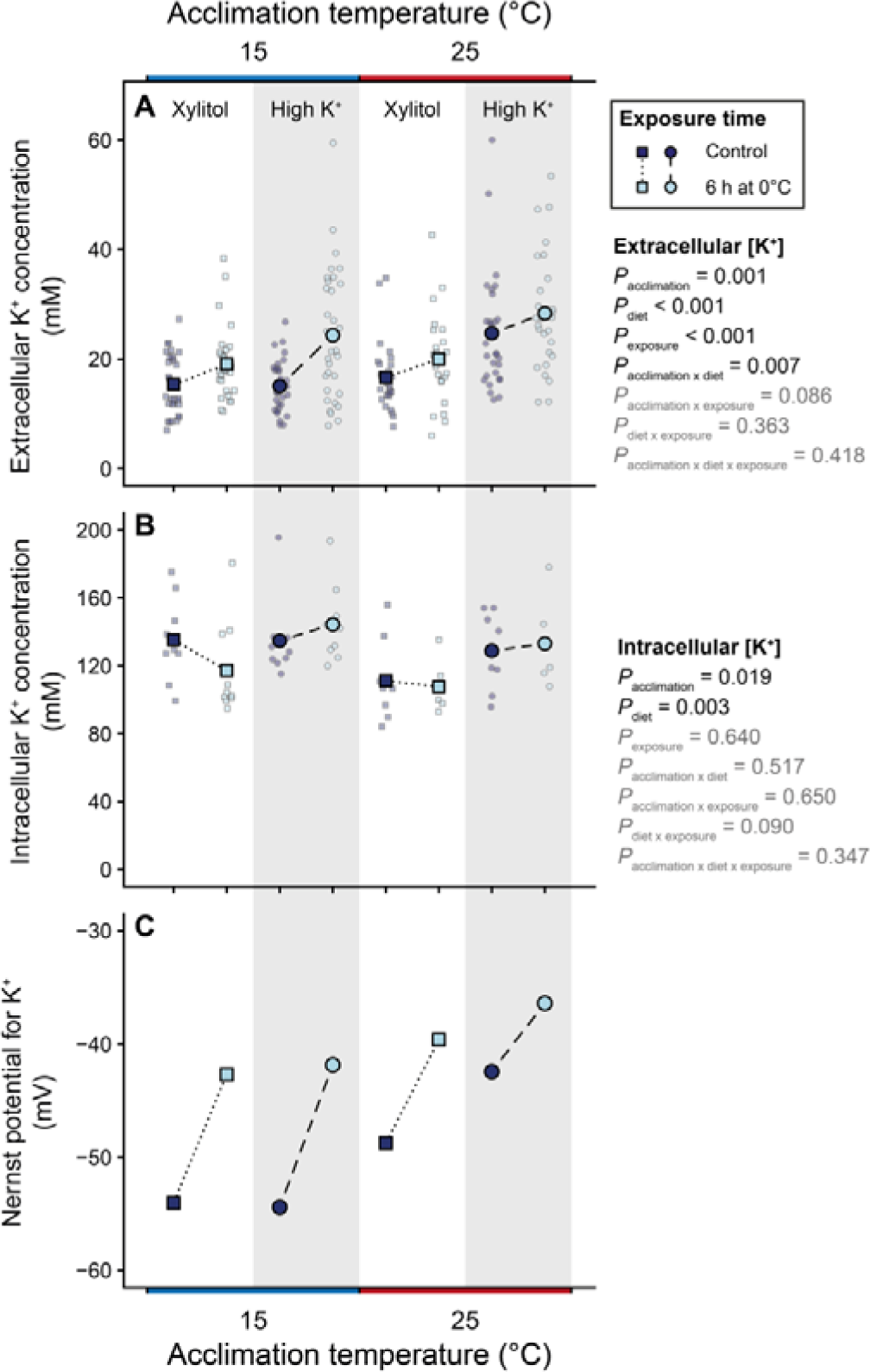
Cold-acclimation mitigates the negative effects of a high-K^+^ diet on K^+^ balance regulation during cold exposure in fruit flies. (A) Extracellular and (B) intracellular [K^+^] was measured in the hemolymph and muscle, respectively, of cold- and warm-acclimated *D. melanogaster* fed either control- (squares, white background) or high-K^+^ (circles, shaded background) diets before (dark blue) and after (light blue) exposure to 0°C for 6 h (N = 22-33 for panel A and 5-10 for panel B). (C) The Nernst potential for K^+^ was calculated for each acclimation, diet, and exposure group based on mean values. For panels A and B the large, opaque points connected with lines represent mean values and smaller translucent points represent individual data points. See **Table 2** for exact, group-specific sample sizes.

Extracellular [K^+^] was measured in the hemolymph, and was ∼ 15 mM in warm-acclimated flies fed the control xylitol diet before being exposed to the cold; however, this was elevated to ∼ 25 mM upon being fed a high-K^+^ diet (effect of diet: F_1,220_ = 18.8, P < 0.001) (**Fig. 2A**). At the same time, there was a significant effect of acclimation temperature (effect of acclimation: F_1,222_ = 12.9, P < 0.001) which interacted with the diet (diet x acclimation interaction: F_1,220_ = 7.3, P = 0.007), such that cold-acclimated flies maintained their extracellular [K^+^] at ∼ 15 mM on both the xylitol-control and high-K^+^ diet. When exposed to 0°C for 6 h, however, the extracellular [K^+^] increased in all treatment groups (effect of exposure: F_1,220_ = 21.8, P < 0.001). This increase in extracellular [K^+^] was the same across diets (diet x exposure interaction: F_1,220_ = 0.8, P = 0.363), but tended to be larger in the cold-acclimated flies (acclimation x exposure interaction: F_1,220_ = 3.0, P = 0.086). No interaction was found between all three factors (acclimation x diet x exposure interaction: F_1,220_ = 0.7, P = 0.418).

Intracellular [K^+^] was measured in thorax (mainly muscle), and was ∼ 111 mM in warm-acclimated flies fed the control xylitol diet before being exposed to the cold; however, this increased by the additive effects of cold acclimation (effect of acclimation: F_1,58_ = 5.8, P = 0.019) and diet (effect of diet: F_1,58_ = 9.9, P = 0.003) resulting in warm-acclimated flied fed the high-K^+^ diet having ∼ 128 mM of K^+^ in their intracellular space, and cold-acclimated flies having ∼ 135 mM intracellular [K^+^] regardless of the diet (acclimation x diet interaction: F_1,58_ = 0.4, P = 0.517) (**Fig. 2B**). A 6 h exposure to 0°C had a tendency to decrease intracellular [K^+^] in control flies with no effect in high-K^+^ fed flies but no effect of exposure reached the level of statistical significance (effect of exposure: F_1,58_ = 0.2, P = 0.640; acclimation x exposure interaction: F_1,58_ = 0.2, P = 0.650; diet x exposure interaction: F_1,58_ = 3.0, P = 0.090). No interaction was found between all three factors (acclimation x diet x exposure interaction: F_1,58_ = 0.9, P = 0.347).

Using the extra- and intracellular concentration of K^+^, it was possible calculate the Nernst potential based on mean values (**Fig. 2C**). Here, warm-acclimated flies fed the control xylitol diet had a Nernst potential of c. −49 mV which was depolarized to c. −42 mV after being fed on the high-K^+^ diet. Conversely, cold-acclimated flies were able to keep their K^+^ Nernst potential stable at c. −54 mV regardless of their diet. Exposure to 0°C for 6 h depolarized the Nernst potential in all treatment groups, and the magnitude of this depolarization was largest in cold-acclimated flies (∼ 12-14 mV) who ended at c. −42 and c. −43 mV for control- and high-K^+^ diets, respectively. The magnitude of the cold-induced depolarization event was less severe in warm-acclimated flies (∼ 6-9 mV), but as their baseline potential was less polarized, their Nernst potential ended up less polarized after cold stress (c. −40 and c. −36 mV for the control- and high-K^+^ diet, respectively).

Intracellular [Na^+^] was measured in conjunction with intracellular [K^+^] and was estimated to be ∼ 43 mM in warm-acclimated flies fed the control xylitol diet (**Fig. 3**). This was value remained unaltered regardless of acclimation regime and diet (effect of acclimation = F_1,58_ = 1.9, P = 0.176; effect of diet: F_1,58_ = 0.1, P = 0.726; acclimation x diet interaction: F_1,58_ = 0.6, P = 0.440), but was substantially reduced in response to the 6 h exposure to 0°C, resulting in all groups reaching intracellular [Na^+^] of ∼ 31-39 mM (effect of exposure: F_1,58_ = 43.4, P < 0.001). Despite this, there was a tendency for this reduction being mitigated in the warm-acclimated flies fed the control diet as this group only experienced a ∼ 4 mM decrease (acclimation x exposure interaction: F_1,58_ = 3.6, P = 0.064). No interaction was found between diet and exposure to cold (diet x exposure interaction: F_1,58_ = 0.1, P = 0.822), nor between all three factors of the experimental design (acclimation x diet x exposure interaction: F_1,58_ = 1.7, P = 0.198).

**Figure 3.**
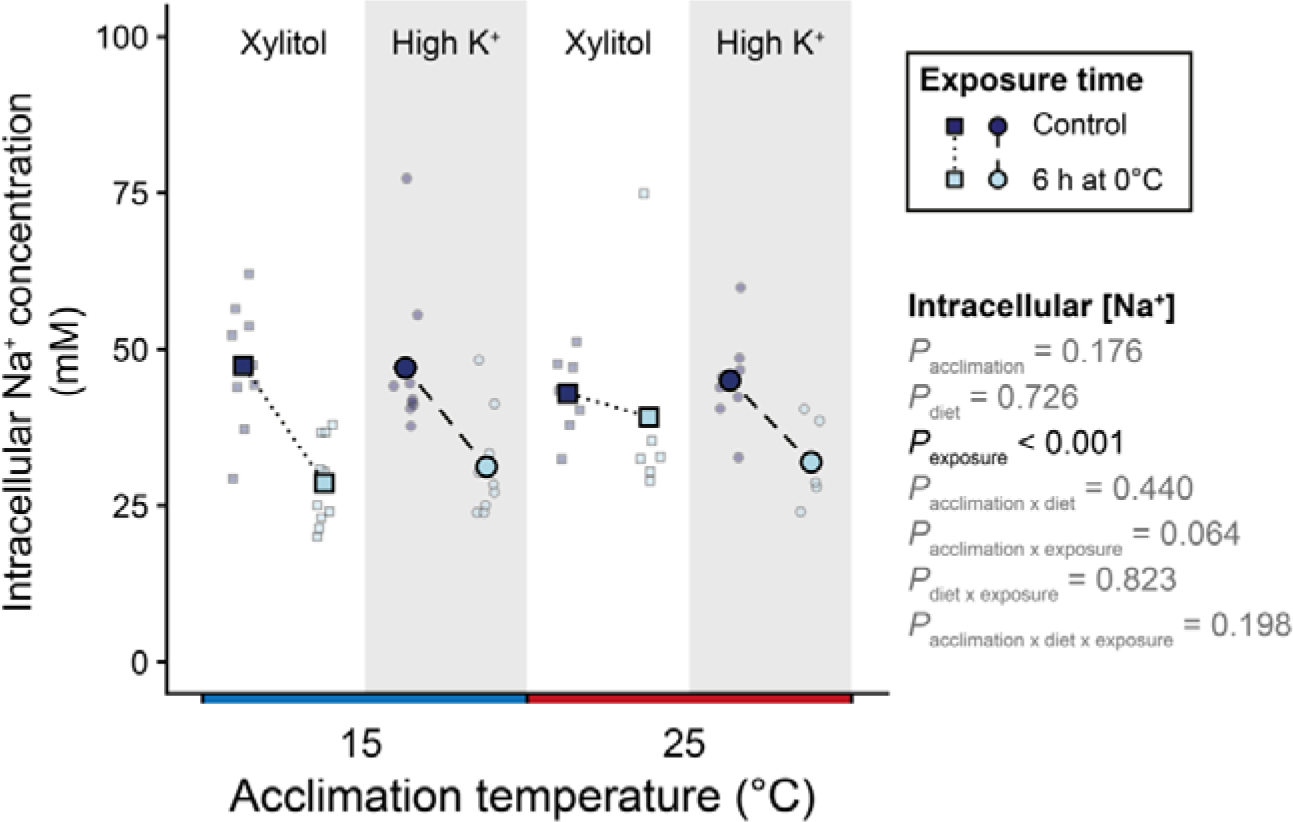
Cold exposure causes a reduction in intracellular Na^+^ concentration in fruit flies. Intracellular [Na^+^] was measured at the same time as intracellular [K^+^] and was therefore measured in the same cold- and warm-acclimated *D. melanogaster* fed either control- (squares, white background) or high-K^+^ (circles, shaded background) diets before (light blue) and after (dark blue) exposure to 0°C for 6 h (N = 5-10). Extracellular [Na^+^] was not measured, and it was therefore not possible to calculate the resulting Nernst potential. Large, opaque points connected with lines represent mean values and smaller translucent points represent individual data points. See **Table 2** for exact, group-specific sample sizes.

## Discussion

Insect performance is directly linked to the environmental temperature, and thermal limits are regularly highlighted as a key driver of species distribution and abundance (Addo-Bediako et al., 2000; Sunday et al., 2019). Thus, there is great interest in elucidating the physiological processes limiting or promoting thermal tolerance such that current distribution limits and range shifts driven by climate change can be understood (Lancaster, 2016). In the present study, we highlight the importance of both thermal and dietary history in insect thermal tolerance by showing that low temperature acclimation and a high-K^+^ diet promote cold tolerance in *Drosophila melanogaster*. We link the increased cold tolerance to a well-established, integrative model for insect cold tolerance (i.e. ‘ionoregulatory collapse’, see MacMillan (2019)), and further speculate that a hitherto unexplored K^+^-handling mechanism might be involved. Lastly, we highlight the need for further cross tolerance research, as these vastly different abiotic stressors (i.e. cold and excess K^+^ load) appear to additively increase cold tolerance through shared mechanisms.

Previous studies have demonstrated that organismal cold tolerance can be altered by feeding on diets with an increased osmolality or salinity, or feeding altogether (Andersen et al., 2013; Yerushalmi et al., 2016; Lebenzon et al., 2020). In the present study we found conflicting effects of feeding salt- or osmolyte-spiked diets on the ability of adult *D. melanogaster* to recover from a cold stress. Specifically, we first found no effect of any dietary modifications on chill coma recovery times (Fig. 1A). However, due to the clear effects of feeding on the high-K^+^ diet, particularly, observed by Yerushalmi et al. (2016), we decided to move forward with the xylitol- and K^+^ diets in the acclimation experiment, where we then found that feeding on a high-K^+^ diet improved both recovery times and survival after cold exposure (see **Fig. 1**). Given that these experiments were performed using the same procedure, on flies of similar age, and by the same experimenters, we are unable to explain these differences in cold tolerance phenotypes. Our diets were similar to those of Yerushalmi et al. (2016), who found improved recovery times (as we did in the second experiment), but no improvements to cold survival despite using the same line of *D. melanogaster*. These seemingly unsystematic differences in experimental outcomes, despite following identical protocols make interpreting the results difficult. However, the most consistent result between studies and experiments is the improved recovery time of the fruit flies after feeding on the high-K^+^ diet when compared to the osmolality-control-fed (xylitol) controls. Interestingly, this goes against the prolonged recovery times elicited by K^+^-feeding in locusts (Andersen et al., 2013), highlighting that the effects of K^+^-feeding on cold tolerance are highly variable and perhaps species-specific. In fact, it can also vary drastically within species, in a genotype-specific manner (Littler et al., 2021). Salt stress tolerance is elicited by a vast variety of physiological and cellular mechanisms, however, central to the response are the Malpighian tubules which act to secrete excess salts, primarily K^+^ for drosophilids (Davies et al., 2014). Thus, it is not surprising that feeding on a high-K^+^ diet might promote K^+^ secretion, which in turn would at least partially aid to prevent hemolymph hyperkalemia during cold exposure. That being said, further research is needed to elucidate the exact molecular, cellular, physiological mechanisms of how high-K^+^ diets improve insect cold tolerance.

Improvements of cold tolerance by the process of low temperature acclimation is well-established for virtually all insect orders and cold tolerance phenotypes (Sinclair et al., 2015; Overgaard and MacMillan, 2017; Weaving et al., 2022). Thus, it comes as no surprise that cold-acclimated *D. melanogaster* were able to recover faster after a cold stress compared to their warm-acclimated conspecifics and were less injured following recovery from a more extensive cold exposure (**Fig. 1**). The mechanisms that promote the improved cold tolerance of cold-acclimated insects has been heavily investigated over the last two decades and relates to multiple physiological processes ranging from 1) improved oxidative phosphorylation and metabolic control (Colinet et al., 2017; Lebenzon et al., 2023), 2) augmented transport capacity in ionoregulatory epithelia (Yerushalmi et al., 2018; Overgaard et al., 2021; Robertson et al., 2023), 3) alteration to the heat shock response (Colinet and Hoffmann, 2012), 4) changes to the immune system (Ferguson et al., 2016; El-Saadi et al., 2023), and 5) overall transcriptomic, metabolomic, and proteomic rewiring (Colinet et al., 2013; MacMillan et al., 2016).

The combined effects of cold acclimation and increased dietary K^+^ loading has not been investigated previously, however, the importance of cross-tolerance between stressors is well-established (Sinclair et al., 2013; Lubawy et al., 2020; Boardman, 2024). In support of our hypothesis, we found that the combination of cold acclimation and a high-K^+^ diet did improve cold tolerance. Specifically, we found a clear additive effect, which improved both recovery times and survival following a stressful cold exposure (see **Fig. 1**C). The physiological mechanisms underlying both improved salt-stress- and cold-tolerance are multiple (see above), however, central to both are changes to the osmoregulatory organs that promote removal of excess K^+^ from the hemolymph: The Malpighian tubules have improved capacity to secrete K^+^ while the hindgut tends to reduce K^+^ reabsorption (Naikkhwah and O’Donnell, 2011; Naikkhwah and O’Donnell, 2012; Davies et al., 2014; Yerushalmi et al., 2018; Andersen and Overgaard, 2020; Overgaard et al., 2021). Thus, our findings here suggest a physiological mechanism related to improved K^+^ handling during cold exposure. Paradoxically, the K^+^ fed flies were generally worse at maintaining hemolymph K^+^ concentrations during cold exposure, despite improved survival. Similarly, only the cold-acclimated group was able to prevent diet-induced hyperkalemia at benign temperatures (**Fig. 2A**). Thus, while eating a high-K^+^ diet disrupts overall K^+^-handling capacity, this is counteracted by cold acclimation. We speculate that the improved cold tolerance despite reduced K^+^-handling capacity could be partially due to a reduced sensitivity to hemolymph hyperkalemia, as observed for cold-acclimated locust muscle previously (Andersen et al., 2017a; Bayley et al., 2020). The hemolymph hyperkalemia observed for K^+^-fed flies is likely due to K^+^ leakage from the K^+^-loaded gut, which has been argued to be a secondary source of cold-induced hemolymph hyperkalemia and which more cold-tolerant flies are better at preventing (Andersen et al., 2017c; MacMillan et al., 2017b; Overgaard et al., 2021). Indeed, changing the electrochemical gradients of ions across the gut epithelium can result in major changes to cold tolerance, as seen in crickets where feeding on a high-Na^+^ diet improves cold recovery times after brief cold exposures at the cost of reduced survival after more chronic exposures (Lebenzon et al., 2020). Regardless, only the cold-acclimated flies fed a high-K^+^ diet are better at maintaining the electrochemical gradient for K^+^ (i.e. the Nernst potential) across muscle membranes during control and cold exposure conditions (**Fig. 2C**). Improved maintenance of the electrochemical potential for K^+^ across the muscle membrane is a well-established mechanism of enhanced cold tolerance in chill-susceptible insects where the degree of hemolymph hyperkalemia experienced during cold exposure generally dictates rates of survival (see Overgaard and MacMillan (2017), Carrington et al. (2020), and Overgaard et al. (2021)). Avoidance of hemolymph hyperkalemia has also been demonstrated for *D. melanogaster* larvae fed on a high-K^+^ diet chronically (Naikkhwah and O’Donnell, 2011). Thus, a shared physiological response to cold acclimation and dietary K^+^ loading being maintenance of the transmembrane K^+^ gradient seems likely. Indeed, multiple stressors can lead to both additive and synergistic thermal performance responses (Kaunisto et al., 2016). The same is true for secretion at the Malpighian tubules, however, the exact cellular and molecular mechanisms often differ (O’Donnell and Spring, 2000). Interestingly, we found that high-K^+^ fed flies, and those cold-acclimated, had increased intracellular (muscle) K^+^ concentrations (**Fig. 2B**). Insect muscle cells are generally not considered to contribute to organismal osmoregulation; however, being an excitable tissue, muscles tightly regulate transmembrane ion gradients to maintain contractility and function, especially at low temperatures (Hoyle, 1953; Hosler et al., 2000; Findsen et al., 2014; Findsen et al., 2016). K^+^-deficient diets are known to reduce intracellular K^+^ in exchange for Na^+^ in human skeletal muscle (McFarlin et al., 2020), thus, even if we don’t see considerable change to intracellular Na^+^ concentration between diets (**Fig. 3**), the opposite K^+^ handling mechanism seems plausible; that the excess dietary K^+^ load might have caused greater reliance on K^+^ in the intracellular space. A higher intracellular K^+^ concentration would further polarize the negative membrane potential which could directly counteract cold-induced depolarization, or mitigate the effects of cold-induced hemolymph hyperkalemia. Indeed, we see that cold-acclimated flies, in particular, are able to protect their K^+^ Nernst potential when fed a high K^+^ diet (Fig. 2C). Alternatively, the challenge of feeding on the high-K^+^ diet might have dehydrated the muscle cells and thereby increased the intracellular K^+^ concentration. However, we found no differences in intracellular water content (see **Fig. S1**) and no corresponding increase in the intracellular Na^+^ concentration when compared to the osmotic control group fed the xylitol-diet (**Fig. 3**). Ultimately, whether what we are observing is a regulated process or a secondary consequence of the high K^+^ diet remains unknown. Regardless of what the exact mechanism might be, we speculate that modifications to K^+^ handling in insect muscle could represent a novel mechanism that promotes insect cold tolerance by preventing cellular depolarization.

The intracellular Na^+^ concentrations were found to only change in response to cold exposure, with a slight tendency for this reduction being larger in cold-acclimated flies (**Fig. 3**). Previous studies measuring muscle concentrations of Na^+^ in cold-acclimated and cold-exposed insects have reported conflicting findings. Specifically, cold-acclimated locusts tend to have higher intracellular Na^+^ concentrations than their warm-acclimated conspecifics (Andersen et al., 2017a). In fruit flies, however, cold acclimation has previously been reported to have no effect on intracellular Na^+^ concentration (MacMillan et al., 2015b), which we corroborate here. Interestingly, the concentrations we report here are more than twice those reported by MacMillan et al. (2015b). The average water content in our samples was ∼ 66% which is slightly below the expected 70-75% (**Fig. S1**); however, our samples did include the thoracic integument and cuticle, which could partially explain this difference. The central paradigm of the current ionoregulatory collapse model of insect chill tolerance is a cold-induced reduction in hemolymph volume caused by movement of Na^+^ (and Cl^-^) from the hemolymph to the gut, leading to a more concentrated hemolymph K^+^ concentration (Overgaard and MacMillan, 2017; MacMillan, 2019). Interestingly, reversing the gradient for Na^+^ across the gut by feeding on a high-Na^+^ diet results in faster recovery times after brief cold exposures but at the cost of survival during prolonged exposures despite no differences in K^+^ handling ability (Lebenzon et al., 2020). In the present study, we did not measure hemolymph Na^+^ concentrations, however, the concentration is well-known to be higher than that of the gut and muscle, creating inward-directed electrochemical gradients (positive Nernst potentials) (Duchǎteau et al., 1953; Andersen et al., 2017b). Low temperature should cause a reduction Na^+^/K^+^-ATPase activity (Cheslock et al., 2021; Andersen et al., 2022), which is a likely culprit for the cold-induced movement of Na^+^ (and Cl^-^ and water) to the gut and could also lead to an increased muscle concentration over time. Cold exposure can also lead to cell swelling, which could partially explain the reduction in intracellular Na^+^ concentration (Boutilier, 2001); however, we saw no corresponding decrease in intracellular K^+^ concentration (**Fig. 2B**) and no change in muscle water content (**Fig. S1**). Thus, further research is needed to investigate how dietary osmotic challenges, excess dietary K^+^ load, thermal acclimation, and cold exposure affect Na^+^ gradients and handling mechanisms.

## Conclusion

The physiological mechanisms that set thermal limits and drive variation in thermal performance traits are important for predictive modelling of animal distribution and abundance. While we are gaining a deeper understanding of chill tolerance, the mechanisms underlying organismal chill tolerance traits are complicated, and represent an emergent property of temperature effects on multiple interacting organ systems. Here we show for the first time that thermal acclimation and diet act additively to improve the cold tolerance of *D. melanogaster* and show that this effect is at least partially tied to ionoregulatory processes occurring in the muscles. We further link this improved tolerance to a well-established model for insect cold tolerance; the ‘ionoregulatory collapse’ model (Overgaard and MacMillan, 2017; MacMillan, 2019). Specifically, we demonstrate that cold- and K^+^-fed fruit flies have improved cold tolerance through an improved ability to regulate the transmembrane gradient for K^+^ by mitigating extracellular hyperkalemia. We posit that increased intracellular K^+^ might be an unexplored K^+^ handling mechanism that aids in maintenance of K^+^ gradients and by extension membrane potentials, resulting in improved cell survival during cold exposure.

## Supplementary material

**Figure S1.**
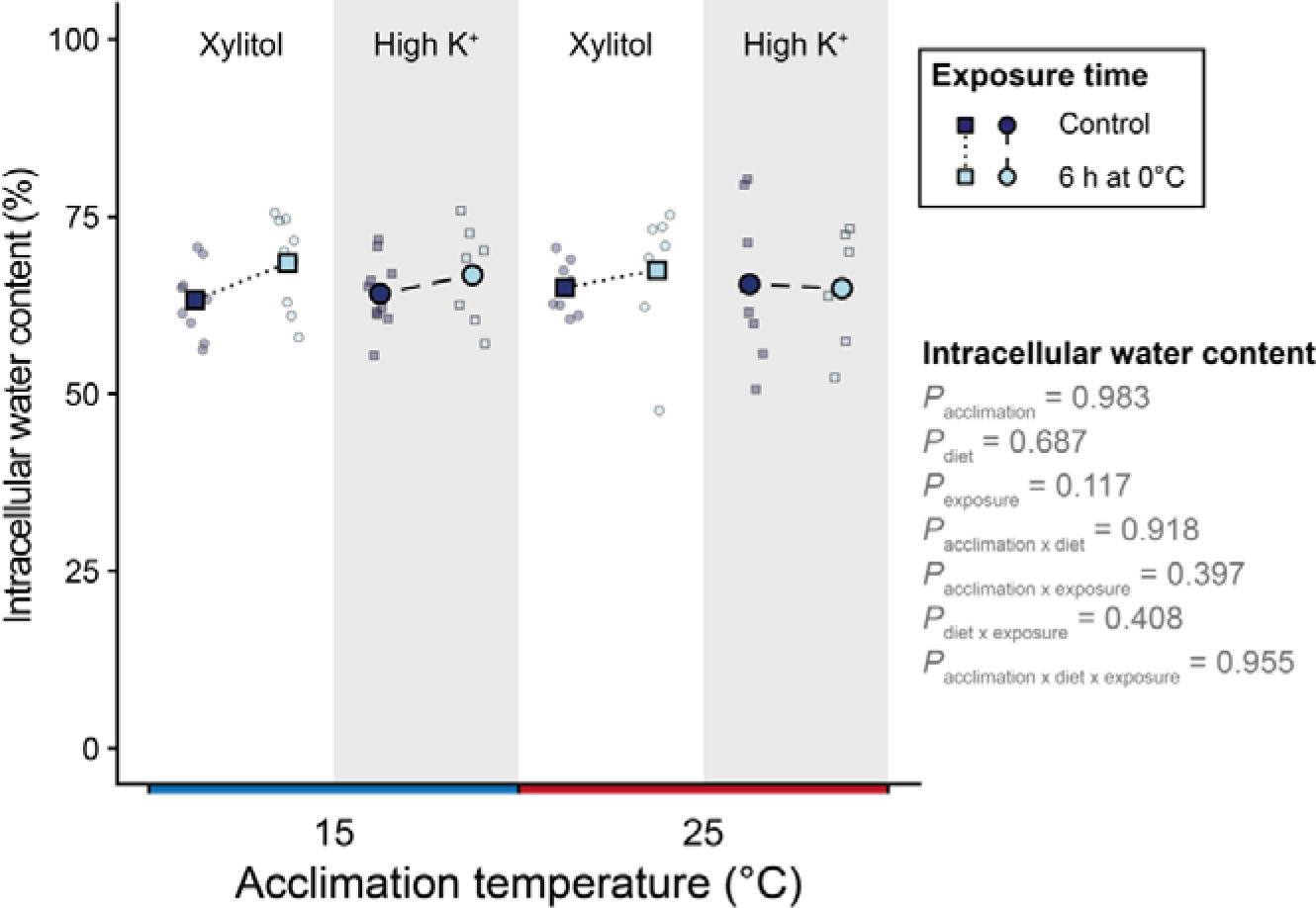
Effects of thermal acclimation, diet, and cold exposure on the intracellular water content measured in *Drosophila melanogaster* thoraces. Total water content was measured during sample preparation for measurement of ion concentrations in thoracic samples, however, here the water content is expressed as a percent of wet weight. As such, the treatments depicted here are the same as those for the intracellular ion concentrations: Cold- and warm-acclimated flies fed either control- (squares, white background) or high-K^+^ (circles, shaded background) diets before (dark blue) and after (light blue) exposure to 0°C for 6 h (N = 5-10). Large, opaque points connected with lines represent mean values and smaller translucent points represent individual data points. See Table 2 for exact, group-specific sample sizes.

